# Global lessons from antibiotic resistance: metformin-hydrolyzing genes in transposable elements, a new threat for type II diabetic patients?

**DOI:** 10.1101/2025.04.28.651035

**Authors:** Tram Vo, Vicky Merhej, Christina Isber, Pierre Pontarotti, Fadi Bittar, Jean-Marc Rolain

## Abstract

Metformin drug, widely used to treat type II diabetic patients, is a major pharmaceutical pollutant of wastewater and rivers. This environmental exposure has driven the evolution of bacteria, such as *Aminobacter* and *Pseudomonas*, to degrade metformin via a Ni²⁺-dependent metformin hydrolase complex (MfmAB). Here we decipher the mechanism of acquisition and horizontal transfer of the *mfmAB* genes, initially mobilized from *Aminobacter* chromosomes to *Pseudomonas* conjugative plasmids via common transposable elements (*IS*1182 and *IS*3/*IS*6 elements) in composite transposons, carrying also other genes involved in guanylurea and biguanides degradation (*guuH* and *bguH*). These mobile elements, historically involved in acquisition of antibiotic-resistant genes from the environment before clonal expansion in clinical settings, now threaten to co-select for both metformin-degrading and antibiotic resistance genes in contaminated waters. This represents a global threat for diabetic patients with concurrent infections that should be urgently added in the roadmap of research in the context of One Health.

## Introduction

The manufacturing and widespread use of pharmaceutical products during the 20^th^ century have contributed to an increasing diversity of active pharmaceutical ingredients and their metabolites in the environment^1^. This extensive release of bioactive compounds into ecosystems can disrupt ecological functions and pose a significant threat to human and ecosystems health^1–3^. Of these compounds, metformin (1,1-dimethylbiguanide), has emerged over the last few decades as a globally dominant pharmaceutical pollutant due to its extensive use and poor biodegradability in wastewater treatment plants (WWTPs) and aquatic environments^1,4–6^.

Originally synthesized in the 1920s, metformin was introduced as a first-line treatment for type II diabetes in France (1959), China (1994), and the USA (1995)^7–9^. Since then, this drug has become one of the most commonly used pharmaceuticals globally, with over 150 million prescriptions worldwide in 2022^10^. More recently, metformin has been investigated for its potential use in the treatment of cancer, endocrine disorders, and obesity, with expectations for its broader use in the future^11^. The direct mode of action of metformin remains incompletely characterized^11^. Evidence now supports the fact that metformin alters the human gut microbiota by mediating its therapeutic effects^12–15^. Intravenous administration of metformin to type II diabetes patients, bypassing the human gut, showed limited to no therapeutic efficacy, further emphasizing the important interaction with gut microbiota in therapeutic effects^16^. After a single oral dose of 0.5 g to 2 g per day, over 80% of metformin is excreted unchanged in urine and feces^17,18^. Due to its limited hepatic metabolism and extensive excretion, metformin has become one of the most predominant micropollutants in WWTPs^1,4^.

Over the past decade, microbial metabolism in the activated sludge of WWTPs has been identified as a major driver of metformin biodegradation^5^. The primary transformation product of metformin in WWTPs, guanylurea, is known to be environmentally toxic, causing reproductive impairment, reduced larval survival, and neurobehavioral alterations in fish after chronic exposure^3^. Bioremediation strategies targeting metformin and its metabolites have therefore been proposed to mitigate environmental contamination. Recent independent studies have identified *Aminobacter* sp. and *Pseudomonas* sp. strains isolated from activated sludge samples in USA, France, and China in 2021 capable of degrading metformin and utilizing it as their sole source of carbon and nitrogen for growth^19–22^. These strains encode a Ni^2+^-dependent heterohexameric enzyme composed of two subunits, MfmA and MfmB, which hydrolyses metformin into guanylurea and dimethylamine^23^. This enzyme belongs to the ureohydrolase superfamily and shares similarity with arginase and agmatinase, highly conserved bacterial enzymes recently identified as metformin targets^18–20,22^. Agmatinase catalyzes the conversion of agmatine into putrescine, a key precursor in polyamine biosynthesis, essential for bacterial growth, nitrogen metabolism, and stress adaptation^18,24^.

Intriguingly, while *mfmAB* genes were reported to be chromosomally encoded in *Aminobacter* spp. strains, they are carried on conjugative plasmids in another *Aminobacter* spp. and in *Pseudomonas* spp. strains in the USA, suggesting their potential horizontal genetic transfer (HGT) ^19–21^. The acquisition of *mfmAB* via mobile genetic elements (MGEs) (i.e conjugative plasmids) could significantly accelerate the global spread of metformin degradation capabilities with implications for both environmental pollution and potentially impacting human metformin treatment. Previous studies have focused mainly on the origin, role and biological activity of this enzyme but the exact genetic support and movement of such enzymes has not been studied so far. This study aims to elucidate the evolutionary origins of the *mfmAB* gene cluster and uncover the genetic mechanisms driving its HGT along with other unexpected genes across bacterial species using advanced computational methods. In the context of One Health, assessing the health risk associated with the spread of such mobile “metformin resistance” genes, by analogy with the current situation of antibiotic resistance emergence and spread in humans, is critical, given the increasing number of patients worldwide being treated with metformin for type II diabetes.

## Materials and methods

### Identification of bacterial genomes containing *mfmAB* and homologous sequences

To investigate the distribution and evolutionary origins of *mfmAB*, the *mfmA* (1,073 bp) and *mfmB* (1,046 bp) sequences from *Aminobacter* sp NyZ550 (NCBI accessions WAX94658.1, WAX94657.1) were used as queries for BLASTn and BLASTp searches against the NCBI non-redundant database in April 2025. To identify potential functional homologs across genera, searches were conducted with a threshold of > 90% identity and > 90% coverage. For each bacterial strain carrying *mfmAB* homologs, relevant metadata were collected, including the geographical location, date, and source of isolation and previously reported metformin- hydrolyzing capacity. In addition, whole-genome phylogenetic analysis of all *Aminobacter* spp. in the NCBI database was performed using TYGS^25^.

### Comparative genomic analysis of metformin-hydrolyzing strains

To elucidate the genetic context and potential transfer mechanisms of *mfmAB* across bacterial species, only genomes of strains with confirmed metformin hydrolase activity were selected for comparative genomic analysis. Plasmid classification and replication origins were determined using MOB-Typer (v.3.0.3) and Ori-Finder, which identify relaxase types and predict plasmid mobility^26,27^. Sequence homology comparisons were performed using the Blastn NCBI tool.

To investigate structural variations and the potential role of MGEs in the horizontal transfer of *mfmAB* across bacterial species, insertion sequences and other MGEs associated with *mfmAB* were identified using ISfinder and mobileOG-db^28^. Genome alignment and synteny analysis were conducted using Easyfig (v.2.2.2) to visualize the genomic context and compare the organization of *mfmAB*-harboring regions across different bacterial genomes^29^.

Illustrations in this manuscript were initially created using BioRender (https://www.biorender.com/) to conceptualize the figures under a full publication license. All figures were subsequently redrawn manually using Microsoft PowerPoint.

## Results

### Genomic identification and evolutionary origin of *mfmAB* in metformin-hydrolyzing strains

BLASTn analysis of *mfmAB* sequences against the NCBI database identified 15 bacterial genomes carrying *mfmAB* homologs with ≥ 99% coverage and 90%-99% identity, including three *Pseudmonas* and 12 *Aminobacter* strains (Table 1). Each genome harbored a single copy of the *mfmAB* homolog within a highly conserved eight-gene cluster spanning approximately 8.2 kb. This cluster encodes a HupE/UreJ family protein, two arginase and agmatinase family proteins (MfmB and MfmA), two hydrogenase nickel incorporation-associated proteins (hypA and hypB), a TetR/AcrR family transcriptional regulator, a cystosine permease (codB), and a second XRE family transcriptional regulator (Fig. 1a). Notably, 11 of the 12 *Aminobacter* strains carried the *mfmAB* homolog on their chromosome, suggesting that the active metformin-hydrolyzing *mfmAB* likely originated from *Aminobacter* spp.

**Fig. 1.**
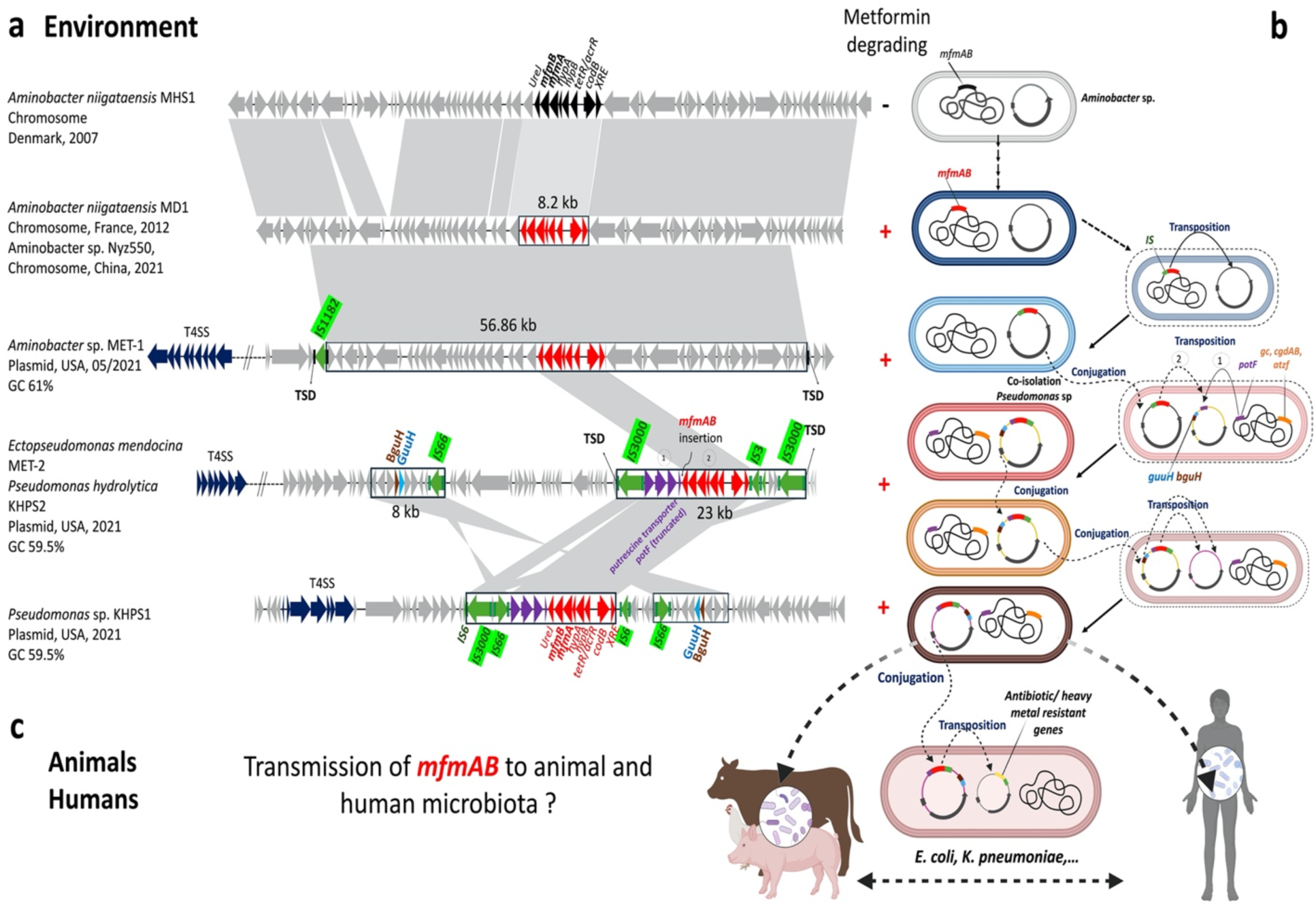
Genomic reconstruction and proposed evolutionary scenario for the acquisition and mobilization of the *mfmAB* gene cluster. a, Comparative genomic alignment of *mfmAB*- containing regions using Easyfig. Coding sequences (CDSs) are shown as arrows: *mfmAB* cluster (red), homologs (black), insertion sequences (green), putrescine transporter genes (violet), biguanide aminohydrolase (brown), guanylurea hydrolase (blue), and type IV secretion system (T4SS) genes (dark blue). Gray shading (81–100% BLASTn identity) indicates conserved regions. **b,** Hypothetical model of *mfmAB* horizontal gene transfer (HGT) and evolution in *Aminobacter* and *Pseudomonas* spp. Chromosomes (black lines) and plasmids (circles) are illustrated with CDSs represented as thick lines and color-coded as in panel **a**. Dashed arrows and circles indicate potential intermediate steps, while solid arrows point post-HGT strains. **c,** Proposed dissemination of *mfmAB* to animal and human microbiota via HGT, driven by co-selection with antibiotic/heavy metal resistance genes (yellow). Gene transfer is suggested to occur via conjugation and transposition. Created in BioRender. Bittar, F. (2025) https://BioRender.com/utwl1xw.

**Table 1.** *mfmAB* homologs identified by BLASTn analysis, detailing genomic localization, strain origins, and validation of metformin-hydrolyzing activity.

| BLASTn alignment results of the *mfmAB* against NCBI Refseq (E-value <1e-10) |  |  |  |  | Strain origins and isolation timeline |  | Functional characterization of MfmAB activity |  |
| --- | --- | --- | --- | --- | --- | --- | --- | --- |
| Organism | Percent Identity | Query coverage | Genome assembly [Accession number hit] | Genomic context (Size in bp) | Source of isolation | Country (region/state)-Year of isolation | Metformin degradation | References |
| *Aminobacter* sp. NyZ550 | 100.00% | 100% | GCA_025628925 [CP114190] | Chromosome (4,947,010) | WWTPs | China-2021 | Confirmed | 22 |
| *Aminobacter niigataensis* MD1 | 99.91% | 100% | GCA_946995915 [NZ_OX341517] | Chromosome (5,500,801) | WWTPs | France (Strasbourg)-2012 | Confirmed | 21 |
| *Aminobacter* sp. MET-1 | 99.86% | 100% | GCA_026521175<sup>a</sup> [JAMSHL010000190] | Plasmid pMET-1 (160,141) | WWTPs | USA (Minnesota)-2021 | Confirmed | 20 |
| *Ectopseudomonas mendocina* MET-2 | 99.63% | 100% | GCA_026503285 [CP098607] | Plasmid pMET-2 (90,221) | WWTPs | USA (Minnesota)-2021 | Confirmed | 20 |
| *Ectopseudomonas hydrolytica* KHPS2 | 99.49% | 100% | GCA_024205225 [CP100554] | Plasmid pKHPS2 (83,915) | WWTPs | USA (Minnesota)-2021 | Confirmed | 20 |
| *Pseudomonas* sp. KHPS1 | 99.58% | 99% | GCA_024205205 [CP100552] | Plasmid pKHPS1 (79,836) | WWTPs | USA (Minnesota)-2021 | Confirmed | 20 |
| *Aminobacter* sp. MSH1 | 92.41% | 100% | GCF_003071665 [CP026265] | Chromosome (5,301,880) | Soil | Denmark-2007 | Negative | 33 |
| *Aminobacter niigataensis* DSM 7050 | 92.32% | 100% | GCF_014200015<sup>a</sup> [NZ_JACHOT010000005] | Chromosome contig (383,642) | Soil | Japan (Niigata)-1992 | Negative | 33 |
| *Aminobacter* sp. DSM 101952 Root100 | 91.90% | 100% | GCF_001424795<sup>a</sup> [NZ_LMCL01000001] | Chromosome contig (720,165) | Root | Germany-2013 | Negative | 33 |
| *Aminobacter aganoensis* DSM 7051 | 91.71% | 100% | GCF_014206975<sup>a</sup> [NZ_JACHOU010000005] | Chromosome contig (301,578) | Soil | Japan (Niigata)-1992 | Negative | 33 |
| *Aminobacter* sp. P9b | 92.35% | 100% | GCF_045345995 [CP150082] | Chromosome (4,937,309) | Ground water | Germany-2016 | ND | - |
| *Aminobacter* sp. SR38 | 91.56% | 100% | GCF_014843375 [CP062112] | Chromosome (5,667,809) | Soil | France-2000 | ND | - |
| *Aminobacter carboxidus* DSM 1086 | 90.44% | 100% | GCF_014863355<sup>a</sup> [NZ_JACZEP010000010] | Chromosome contig (155,224) | Soil | Russia-1977 | ND | - |
| *Aminobacter anthyllidis* D-10A | 90.24% | 100% | GCF_029872135<sup>a</sup> [NZ_JARXNE010000001] | Chromosome contig (562,047) | Sediment | China-2022 | ND | - |
| *Aminobacter anthyllidis* LMG 26462 | 90.11% | 100% | GCF_018555685<sup>a</sup> [NZ_JAFLWW010000004] | Chromosome contig (670,596) | Root | France-2011 | ND | - |

<sup>a</sup> incomplete genome.
WWTPs, wastewater treatment plants. ND, not determined<sup>33</sup>.

A total of six strains, including three *Aminobacter* spp. (NyZ550, MD1, and MET-1) and three *Pseudomonas* sp. strains (MET-2, KHPS1, and KHPS2), were previously reported to hydrolyze metformin through enzymatic assays^19–22^ (Table 1). These isolates were recovered from WWTPs in France in 2012, in China and the USA in 2021^19–22^. Interestingly, a genome-based phylogenomic analysis indicated that these metformin-hydrolyzing *Aminobacter* strains, isolated from three different continents, belong to three different lineages (Supplementary Fig. 1).

Furthermore, localization analysis of *mfmAB* revealed that in two *Aminobacter* strains (NyZ550 from China and MD1 from France), *mfmAB* was chromosomally encoded, whereas in four strains from the USA (*Aminobacter* MET-1 and three *Pseudomonas* strains), *mfmAB* was located on plasmids, indicating evidence of the HGT of *mfmAB* across bacterial species, particularly in the USA (Table 1).

### Transposition of *mfmAB* from the chromosome to a plasmid of *Aminobacter* sp

The origin of replication *repC* of *Aminobacter* sp. plasmid pMET-1 containing *mfmAB* has 82% identity and 100% coverage with a plasmid from *Rhizobiaceae bacterium* strain isolated in an activated sludge from Hong Kong (GCA_023953835.1). Comparative genomic alignment between pMET-1 and the chromosomes of *Aminobacter* sp. MD1 and NyZ550, revealed a homologous region of approximately 56.86 kb containing *mfmAB* cluster (Fig. 1a). Notably, this region on pMET-1 was flanked by a complete *IS*1182- family insertion sequence (*IS*). Further examination revealed the presence of two inverted repeats left (IRL) and right (IRR) associated with *IS*1182, along with an additional IRL structure located at the end of the homologous region (Fig. 1a). Importantly, a 5 bp-directed repeat sequence indicative of a target site duplication (TSD) was observed surrounding this inserted region, providing strong evidence of a recent transposition of *mfmAB* from the chromosome to plasmid pMET-1 of *Aminobacter* sp. mediated by *IS*1182 (Fig. 1a, b). Together, these findings support that the transposition of the *mfmAB* is rather a recent evolutionary event that is compatible with the recent selective pressure with metformin by human prescription in patients with type II diabetes.

### Horizontal transfer of an *mfmAB*-carrying *Aminobacter* plasmid followed by transposition into *Pseudomonas* plasmids

All four metformin-hydrolyzing strains from Minnesota, USA, harbored an *mfmAB* cluster on plasmids encoding a complete set of conjugation-associated genes. These included a type IV secretion system (T4SS) with *virB4* (pMET-1, pMET-2, pKHPS2) and *tra* cluster genes (pKHPS1), as well as an origin of transfer (*oriT*), relaxase, and transfer coupling protein (Fig. 1a). These genetic features suggest that these plasmids are mobilizable and capable of conjugative transfer. However, comparative plasmid analysis revealed that pMET-1 in *Aminobacter* sp. strain is distinct from the plasmids in the three *Pseudomonas* sp. strains (Fig. 1a). The alignment sequence of pMET-1 with those of *Pseudomonas* sp. strains, identified a conserved 8.2 kb region containing *mfmAB* (Fig. 1a). At the distal end of this region, various complete *IS*s were observed, including an *IS*3 element in pMET-2 and pKHPS2, and an *IS*6 element in pKHPS1 (Fig. 1a). Importantly, each *Pseudomonas* sp. strain harbored only one plasmid, suggesting that the *mfmAB* cluster in three *Pseudomonas* sp. strains was not acquired via direct conjugation and stable maintenance of an *Aminobacter* conjugative plasmid such as pMET-1. Instead, the transfer likely occurred through an initial conjugative transfer of the *mfmAB*-carrying plasmid from *Aminobacter* sp. to the *Pseudomonas* sp. strains, followed by the *IS*-mediated transposition of the *mfmAB* cluster into resident *Pseudomonas* plasmids, with a subsequent loss of the original donor plasmid (Fig. 1b).

Interestingly, in three *Pseudomonas* plasmids, this cluster of 8.2 kb *mfmAB*-containing region was found within a larger 23 kb composite transposon flanked by *IS*3000 elements, belonging to the Tn3-family transposase that contains a cluster of genes encoding polyamine ABC transporter substrate-binding proteins and a partial spermidine/putrescine ABC transporter potF (Fig. 1a). A BLASTn search for these transporter genes revealed high sequence identity (88%) and coverage (98%) with homologous regions in the chromosome of *Pseudomonas* sp. strains, suggesting that this cluster was initially chromosomally encoded before being transposed onto plasmids (Fig. 1b). The truncation of *potF* adjacent to the *mfmAB* cluster indicates that the transposition of *mfmAB cluster*, mediated by either *IS*3 or *IS*6 elements, occurred after the initial acquisition of *Tn*3-family transposon containing these transporter genes (Fig. 1b).

### Plasmid-mediated metformin/biguanide/guanylurea degrading in *Pseudomonas sp.* strains

Comparative sequence analysis of the three *Pseudomonas* plasmids showed that pMET-2 of *Ectopseudomonas mendocina* and pKHPS2 of *Pseudomonas hydrolytica* revealed a high similarity (99.99% identity and 92% coverage), strongly supporting the occurrence of conjugative transfer of *mfmAB*-carrying plasmid between these two *Pseudomonas* sp. strains. In contrast, pKHPS1 exhibited significant genetic divergence but shared two distinct homologous regions with pMET-2 and pKHPS2 (Fig. 1a). In addition to *Tn*3-like transposon containing *mfmAB*, these plasmids harbored a second 8 kb-conserved region, that is flanked by *IS*66, encoding guanylurea hydrolase (*guuH*) and biguanide aminohydrolase (*bguH*), enzymes known to degrade guanylurea and biguanide, respectively^20,30^ (Fig. 1a and Fig. 2). This cluster was rather transposed by the *IS*66 element, known to be a transporter-related insertion sequence between different plasmids of *Pseudomonas* sp. strains^31^ (Fig. 1a, b).

**Fig. 2.**
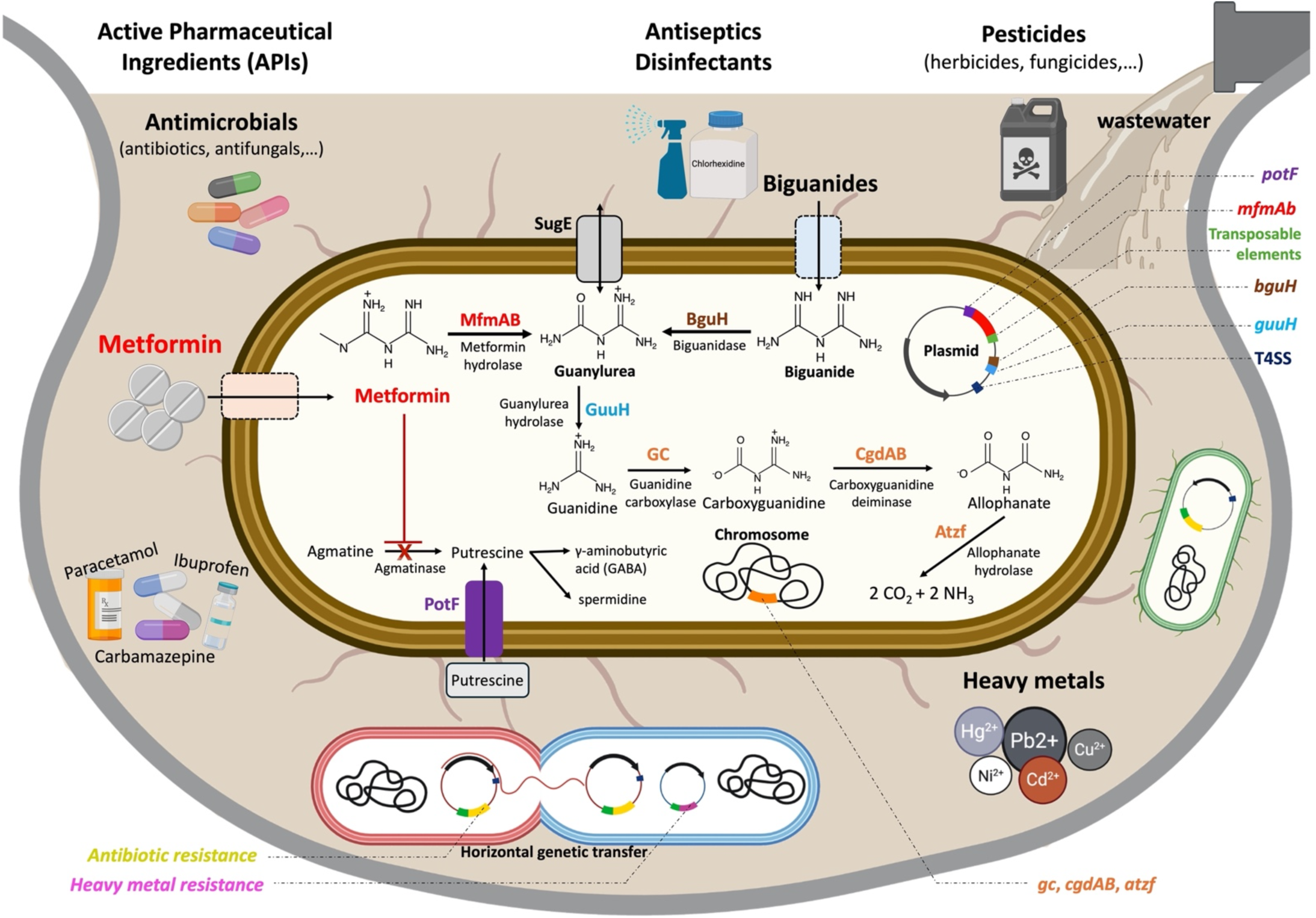
Wastewater environmental microcosm: the PharmacoMicrobioResistance paradigm. Degradation pathways of metformin and biguanide by environmental bacteria. Metformin- hydrolyzing isolates encode MfmAB enzymes (red), the putrescine transporter PotF (violet), biguanide aminohydrolase BguH (brown), and guanylurea hydrolase GuuH (blue), type IV secretion system (T4SS) genes (dark blue), all represented as thick lines. These genes, associated with various transposable elements (green), are located on plasmids (circles). Additional enzymes for complete catabolism of metformin such as guanidine carboxylase (GC), carboxyguanidine deiminase (CgdAB), and allophanate hydrolase (AtzF) which are shown as thick orange lines and encoded on the chromosome (black lines). Transporters for guanylurea SugE (grey), metformin (light orange) and biguanides (light blue) are represented as rectangles. Created in BioRender. Bittar, F. (2025) https://BioRender.com/oaz6y67.

## Discussion

### The Red Queen runs on metformin: plasmid evolution and bacterial adaptation to anthropogenic pollutants

As one of the most widely used medications, metformin has become one of the most frequently detected pharmaceuticals in wastewater, with concentrations reaching up to 16 000 ng/L in the USA^1,4,5,7,20^. This persistent environmental presence is believed to impose selective pressure on microbial communities, facilitating HGT and the evolution of metformin-degrading capabilities, such as those conferred by the *mfmAB* gene cluster^32^ (Fig. 1a, b and Fig. 2). The recent and almost concomitant discovery of environmental bacteria capable of hydrolyzing metformin on three different continents (China, Europe, and the America) is intriguing and we hypothesized that this was not a coincidence but rather a widespread phenomenon on a global scale due to the recent epidemic of diabetic patients and the massive pollution of water environments by this drug (Fig 2). Previous studies hypothesized a chromosomal origin for *mfmAB* gene cluster in *Aminobacter* in the environment, none comprehensively characterized its genetic context or clarify the precise mechanism/scenario of its mobilization from the chromosome of *Aminobacter* to various plasmids using advanced computational methods^20,23,33^. Here we confirm the *Aminobacter* chromosomal origin of these genes and we show that these genes evolved convergently in three independent lineages (Asia, Europe, and North America), supporting Darwinian principles of adaptation (Fig. 1a, b and Supplementary Fig. 1). A precise comparative analysis of the available genomes allowed us to reconstruct the scenario of mobilization of this cluster in the bacterial species analyzed (Fig. 1b). The cluster was firstly mobilized from the chromosome to a conjugative plasmid in *Aminobacter* by transposition via a common *IS*1182. Conjugation of this plasmid into *Pseudomonas* resulted in this transposon jumping to another *Pseudomonas* conjugative plasmid, on at least two independent occasions via an *IS*3 or an *IS*6 element, into larger composite transposons that already contain an additional cluster of four genes associated with putrescine uptake^34,35^ resulting in the truncation of adjacent genes (Fig. 1a, b). Finally, a second 8 kb-cluster encoding for guanylurea hydrolase (*guuH*) and biguanide aminohydrolase (*bguH*), enzymes known to degrade guanylurea and biguanide, was also transposed in these plasmids using an *IS*66 element^20,30^ (Fig. 1a, b). This work highlights the very recent and dynamic process of evolution of these plasmids in the context of environmental pollution by metformin and the constant and formidable adaptation of bacteria to survive in rapidly changing environments polluted by anthropogenic activities. This evolutionary trajectory aligns with the Red Queen hypothesis, which suggests that organisms must continuously evolve not just to gain advantage, but simply to survive under environmental pressures. Although environmental pollution by metformin has persisted for decades, metformin hydrolase was discovered only recently in 2021^1,4–6^. This suggests that bacteria initially endured metformin exposure without this enzyme, likely relying on alternative resistance mechanisms. The transposable elements characterized in our study may represent a key evolutionary adaptation, enabling bacterial survival under prolonged metformin selection pressure. Since metformin inhibits agmatinase, which is a key enzyme to produce putrescine, bacteria were able to counteract this inhibition by expressing a putrescine uptake transporter, enabling direct environmental acquisition of this nutrient^18,34^ (Fig. 2). The transposition of the *mfmAB* cluster into this initial transposon and truncation of this transporter was not deleterious for the bacteria because the hydrolysis of metformin restores the biosynthesis of putrescine by the agmatinase. However, metformin hydrolase produces guanylurea that is thought to be a toxic compound for bacteria and detoxification of this compound could take place due to the guanylurea hydrolase *guuH* present in the second transposable element that makes it possible to further use guanylurea as a source of carbon and nitrogen^20^ (Fig. 2). Once again, transposition proves critical for bacterial survival and adaptation to shifting nutrient conditions.

### Natural occurrence of metformin analogs and global lessons from antibiotic resistance dissemination

Metformin is an alkaloid biguanide with two methyl substituents, forming dimethylbiguanide. It is derived from galegine, a natural analog of metformin isolated from the traditional herb *Galega officinalis*^7,8^. The synthesis of metformin simply relies on the ability of arginine to transfer its amidine group to a precursor of galegine by a transamidination reaction^36^. Interestingly, a vast selection of guanidine derivatives can be found in nature and may explain the ancient origin of the “metformin hydrolase” cluster^8,36^. Indeed, secondary metabolites bearing a guanidine moiety have been isolated from several bacterial species such as *Streptomyces* spp., *Amycolatopsis orientalis*, and *Xenorhabdus* strains^36^. Non-ribosomal peptides, many of which with agmatine or a modified arginine residue have been found in *Cyanobacteria* sp., such as *Microcystis* sp^36^. Guanidine derivates have also been isolated from the marine sponge Crambe crambe and plants such as *Solanum cernuum* and marine nudibranch *Actinocyclus papillatus* (Mollusca)^36^. An imidazole-bis-guanidine alkaloid has been isolated from zoanthid *Epizoanthus illoricatus*^36^. The structural similarity, leading to prolonged bacterial exposure to guanidine analogs in natural environments, may have driven the evolutionary transfer of metformin- hydrolyzing enzymes. Furthermore, the widespread presence of these analogs could explain the independent emergence of an arginase–agmatinase-related gene pair identified in *Aminobacter* strains from three different continents, likely as a result of convergent evolution^19–21^. The widespread distribution of agmatinase family enzymes across diverse bacterial phyla suggests an ancient evolutionary connection between bacterial metabolisms and guanidine derivatives^23^.

Notably, the metformin metabolism may have originated from ancestral pathways linked to agmatinase activity^18,37^. The agmatinase homologs might have played a potential role as a reservoir for novel metformin-metabolizing enzymes. The evolutionary scenario of *mfmAB* in metformin- hydrolyzing strains can be conceptualized through a framework that mirrors the emergence and dissemination of antimicrobial resistance genes^38–42^ (Fig. 2). The recently described plasmid- mediated mobile colistin-resistant (*mcr-*1) gene that threatens the clinical efficacy of colistin in patients infected by multidrug-resistant bacteria is the best example of the evolution of an environmental ancestral gene to a specialized enzyme mobilized via transposition in clinical bacteria within a very efficient and rapid turnaround time^38,43^. Since its first report in *Escherichia coli* isolates from pigs in China, the *mcr*-1 gene has been isolated worldwide in humans and animal reservoirs, and in environmental sources such as wastewater, soil, and rivers^38,44,45^. The widespread use of colistin in agriculture has facilitated the rapid and uncontrolled dissemination of plasmid- mediated *mcr*-1 genes among bacteria^43^. Global lessons from this example have shown that the ancestor of this gene has evolved from different bacterial species in the aquatic environment as a convergent evolution according to Darwin’s theory before being mobilized via transposition events in conjugative plasmids in animals. Colistin selective pressure in animal husbandry then amplified the spread in animals that later transferred to clinically relevant bacteria, and that is now pandemic^46^. A similar concern is emerging with antifungal agents, as the extensive agricultural use of azole-based fungicides is now a major driver of resistance in *Aspergillus* species and probably the emergence of *Candida auris*^47^. Here we show that various *IS* elements and transposition events from the chromosome of an environmental progenitor to conjugative plasmids of metformin hydrolases are exactly the same as those recently described for the antibiotic resistance global crisis^42,48^. Indeed, *IS*1182, *IS*3 and *IS*6 are the most frequent mechanisms involved in the transposition of current pandemic antibiotic-resistant genes (vancomycin resistance operon *vanA*, *mcr-1*, extended-spectrum beta-lactamases, carbapenemases, aminoglycosides modifying enzymes, etc.) frequently organized into plenty multiple different multidrug-resistant composite transposons within various conjugative plasmids^41,42,48–50^. The high mobility and rapidity with which these MGEs reorganize themselves according to the selection pressures exerted in the ecosystems in which bacteria evolve, in particular anthropogenic environments polluted simultaneously by antibiotics and metformin and its metabolites, pose a major risk to humanity of transfer of this transposon of enzymatic metformin “resistance” genes onto multidrug-resistant transposons and/or conjugative plasmids in clinical bacteria (Fig. 1c).

### Potential clinical impact of metformin-hydrolyzing enzymes and future therapeutic strategies

To date, there is no evidence of the presence of this transposon in the human or animal digestive microbiota, which means that transfer from the environment has not yet taken place or has not yet been reported because it has not been specifically searched for. However, clinical reports of metformin treatment failure in patients with type II diabetes along with the expected increase in diabetic patients in the years to come raises concerns about the possible HGT of *mfmAB* from environmental bacteria to human-associated microbes potentially compromising the therapeutic efficacy of metformin^38,40,51^ (Fig. 1c). Interestingly, it has been recently discovered that specific enzymes encoded by the human gut and oral microbiome can inactivate another antidiabetic drug, acarbose, through phosphorylation, suggesting a specific form of microbiome-mediated resistance that may impact both microbial competition and the drug’s clinical effectiveness^52^. Global lessons from antibiotic resistance may be helpful to start designing clinical studies to look for potential synergistic effect of metformin. Notably, sulfonylurea agents such as gliclazide, which also exhibit antimicrobial properties, have demonstrated therapeutic benefits when used in combination with metformin in patients with suboptimal glycemic control compared to metformin alone^53^. Similarly, we believe that it is also timely to encourage research looking for metforminase inhibitors similarly to beta-lactamase inhibitors. Metformin hydrolase is a Ni2+ dependent enzyme that belongs to a large superfamily of metalloproteins with diverse biological functions distributed in all domains of life including archaea, bacteria, CPR, and humans^54^. In the context of antibiotic resistance, the development of metallobetalactamase inhibitors remains challenging, however novel boronate-based inhibitors such as taniborbactam was shown to be effective against such enzymes^55^. Collaborative efforts with clinical microbiologists and endocrinologists should be undertaken to promote the development of boronate-based inhibitors and/or other metalloenzyme inhibitors in this context. Similarly, the inactivation of such metalloenzymes by metal chelators that remove nickel ions or metal-based drugs that replace nickel with another metal are other avenues of research that should be considered in the future.

In conclusion, our study shows that currently the mobilization of the transposon carrying *mfmAB* into plasmids of *Aminobacter* and *Pseudomonas* has been reported only in environmental water in Minnesota, USA, suggesting that it would be opportune to develop global strategies and take decisions requiring evidence-based risk assessments. The potential co-selection and formation of composite transposons carrying all metformin-hydrolyzing, antibiotic and heavy metals resistance genes, that could later spread to clinical settings should urgently be monitored and actively surveyed around the world (Fig. 2) ^56,57^. Lessons should be learned from the emergence and spread of antibiotic resistance in the context of One Health to limit such scenario. Finally, many other pharmaceutical agents widely used in humans for different diseases are now known to be major pollutants and to be degraded by bacterial communities in the environment (such as paracetamol, ibuprofen, carbamazepine)^58–60^ (Fig. 2). Therefore, we strongly believe that a new research field, that we propose to call “PharmacoMicrobioResistance”, (i.e. microbial resistance to non-antibiotic human-targeted drugs) should be added, to encompass pharmaceutical agents other than antibiotics in the current research proposals in the context of One Health.

## Declarations

### Ethical approval

Not required.

## Acknowledgements

We are very grateful to the IHU Méditerranée Infection, Marseille, France for financial support.

## Supplementary data

**Supplementary Fig. 1.**
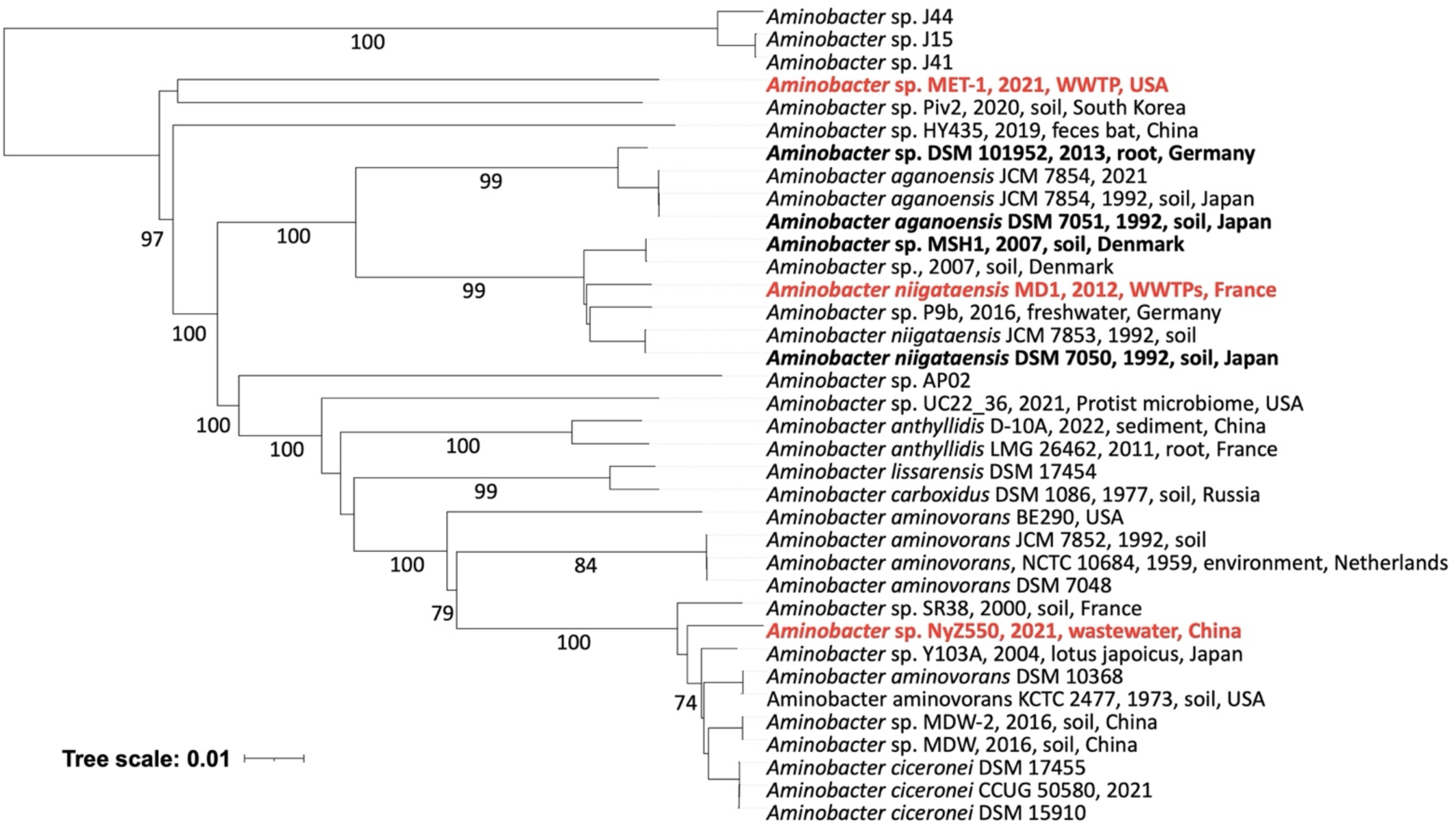
Phylogenetic tree based on whole-genome sequences of *Aminobacter* spp. available in the NCBI database. Strains known to hydrolyze metformin are highlighted in bold red, while non-metformin-hydrolyzing strains are shown in bold black. Other strains, shown in regular font, have not been previously tested for metformin degradation^20,21,33^.

